## Supplemental file for "Molecular mechanics underlying flat-to-round membrane budding in live secretory cells"

### Supplementary Information

Supplementary information contains Supplementary Figs. 1-7 and captions of Movie S1-S8.

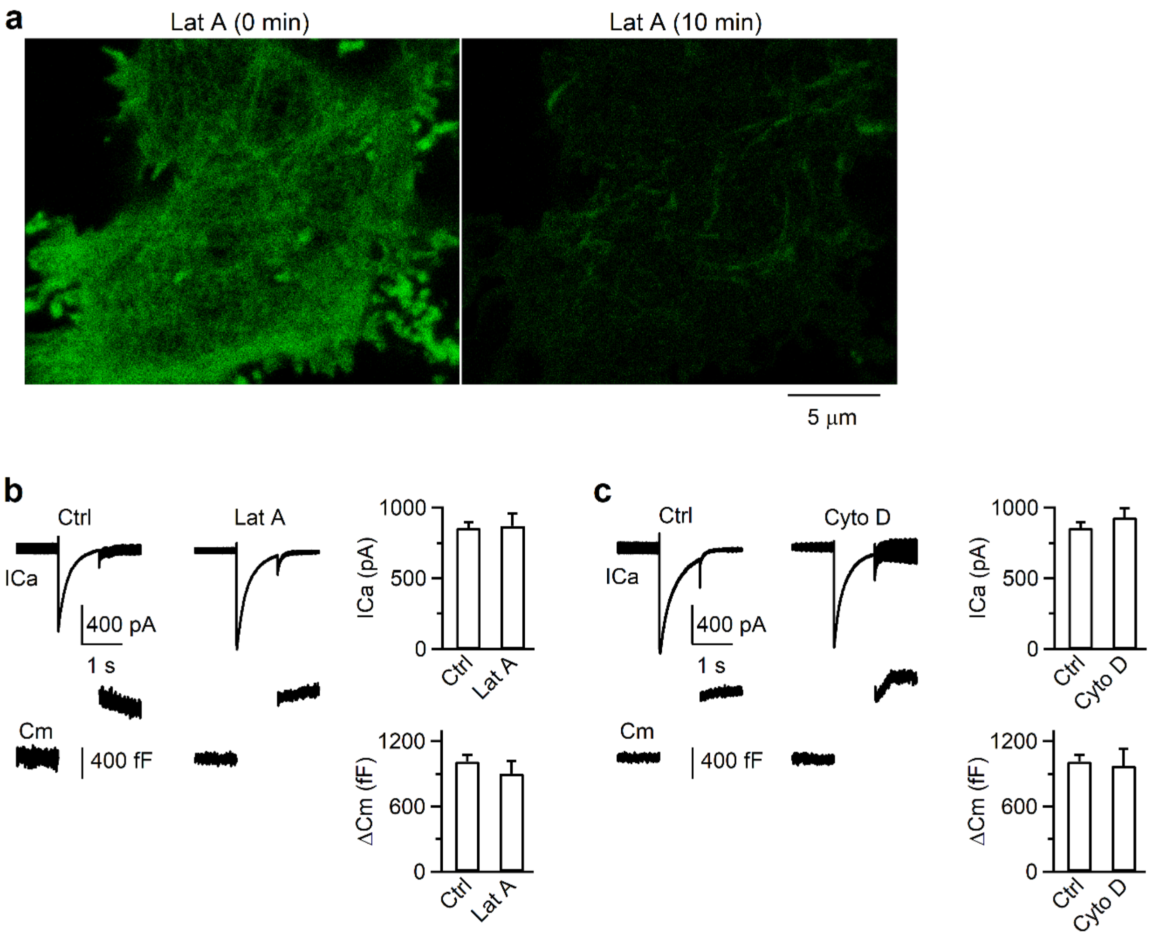


**Supplementary Figure 1. Latruculin A or Cytochalasin D discupts F-actin, but does not affect ICa and capacitance jump.**

(**a**) F-actin labelled with lifeact-mNeonGreen before (0 min) and 10 min after Latruculin A (Lat A, 3 µM) application in a chromaffin cell (confocal images).

(**b**) Left: sampled ICa (upper) and Cm (lower) induced by $depol_{1s}$ in control.
Middle: sampled ICa (upper) and Cm (lower) induced by $depol_{1s}$ in the presence of Lat A (3 µM).
Right: The amplitude (mean + s.e.m.) of ICa (upper) and ΔCm amplitude (lower) induced by $depol_{1s}$ in control (513 cells) and in the presence of Lat A (3 µM, 61 cells).

(**c**) Similar arrangements as in **b**, except in control (513 cells) or in the presence of Cytochalasin D (Cyto D, 4 mM, 57 cells).

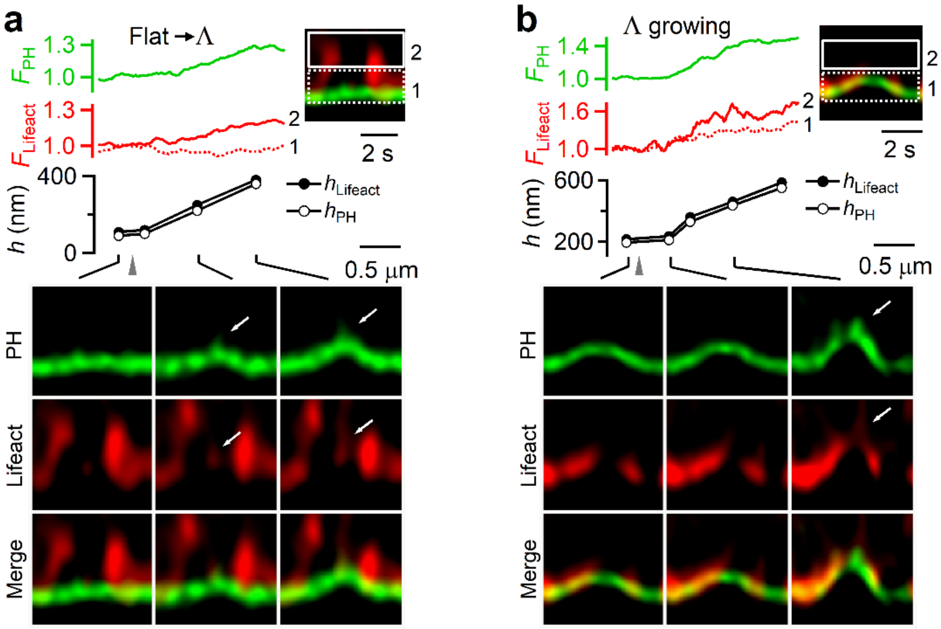


**Supplementary Figure 2. F-actin in the act of pulling membrane inward**

(**a-b**) $F_{PH}$, lifeact-mTFP1 fluorescence ($F_{lifeact}$), $PH_G$-labelled Λ's $h_{PH}$, the highest position of lifeact-mTFP1-labelled F-actin associated with Λ ($h_{lifeact}$), and sample $XZ/Y_{fix}$ images showing F-actin filament recruitment, attachment at, and movement with the growing Λ's tip during Flat→Λ. $F_{lifeact}$ from regions 1 (near Λ's base) and 2 (above Λ's base, inset) are plotted. spike-like protrusion attached to growing F-actin filaments (arrows). $h_{PH}$ and $h_{lifeact}$ were measured as the height from the $PH_G$-labelled base membrane. **a** and **b** show two examples; they are the same as Figs 1f and 1g, respectively, except that lifeact fluorescence is plotted with less saturation. Such a plot allowed us to see strong lifeact signals better, but weak signals worse (e.g., lifeact at Λ's tip).

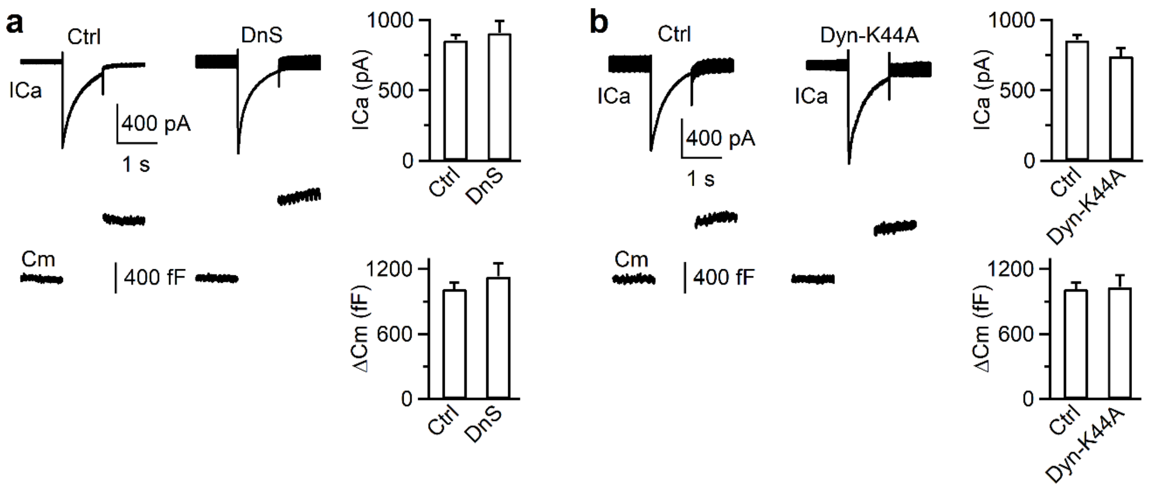


**Supplementary Figure 3. Inhibition of dynamin does not affect calcium currents or capacitance jumps.**

(**a**) Left: sampled ICa (upper) and Cm (lower) induced by $depol_{1s}$ in control.

Middle: sampled ICa (upper) and Cm (lower) induced by $depol_{1s}$ in the presence of dynasore (80 μM).

Right: The amplitude (mean + s.e.m.) of ICa (upper) and ΔCm amplitude (lower) induced by $depol_{1s}$ in control (513 cells) and in the presence of dynasore (DnS, 80 μM, 64 cells).

(**b**) Similar arrangements as in **b**, except in different conditions: without (513 cells) or with overexpression of dynamin 1-K44A (Dyn-K44A, 106 cells).

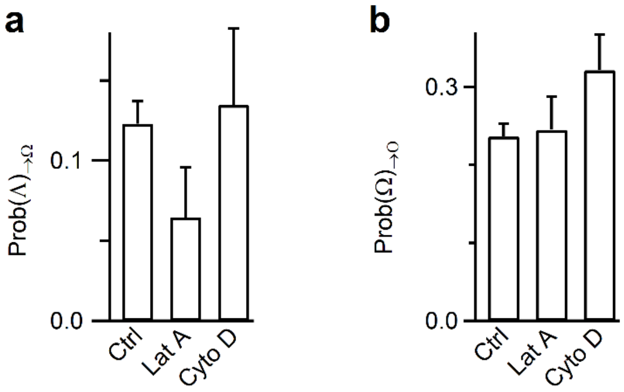


**Supplementary Figure 4. Inhibition of F-action does not significantly affect Λ→Ω or Ω→O transition.**

(**a**) Prob(Λ)$_{\rightarrow\Omega}$ in control (Ctrl, 513 cells), latrunculin A (Lat A, 61 cells), or Cytochalasin D (Cyto D, 57 cells). Data are not significantly different ($p > 0.05$, t test, compared to Ctrl).

(**b**) Prob(Ω)$_{\rightarrow O}$ in control (Ctrl, 513 cells), latrunculin A (Lat A, 61 cells), or Cytochalasin D (Cyto D, 57 cells). Data are not significantly different ($p > 0.05$, t test, compared to Ctrl).

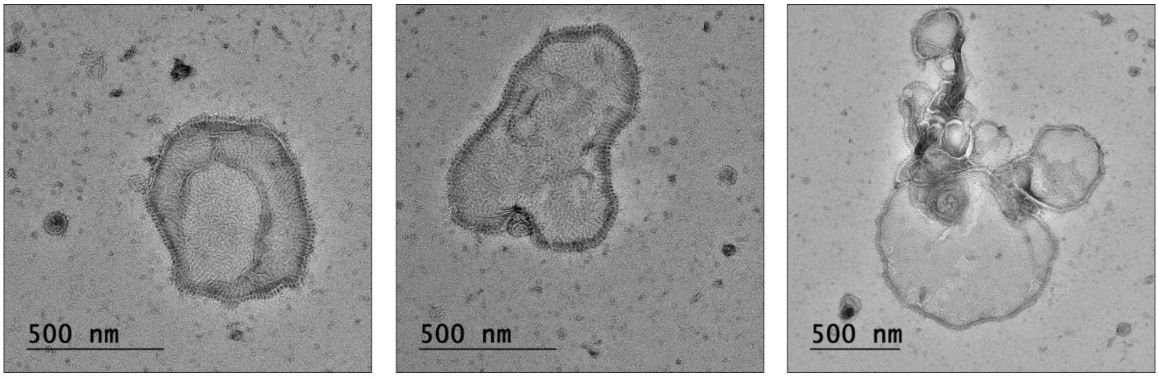


**Supplementary Figure 5. Dynamin decoration of large liposomes**

Sampled negative stain images of ΔPRD-dynamin 1 decorated around large DOPS vesicles after 30 sec of incubation with the lipid.

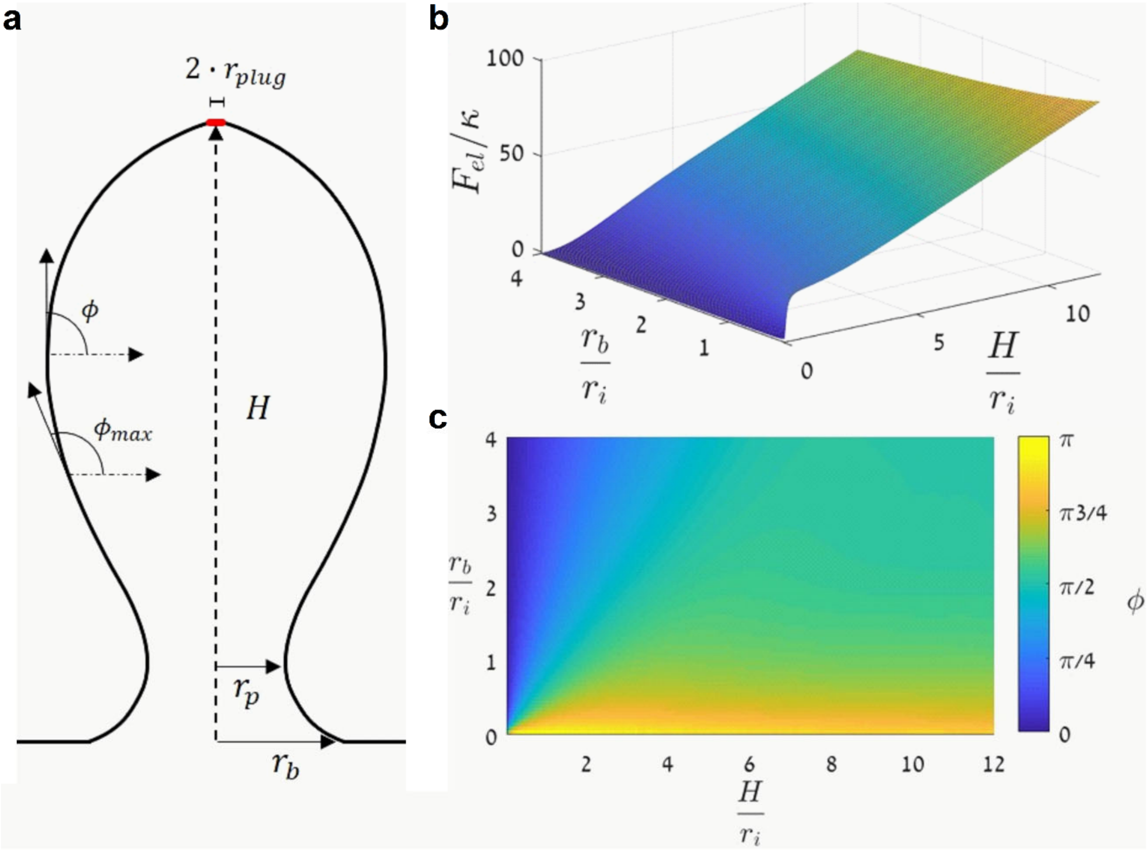


**Supplementary Figure 6. Model and parameter definitions.**

(**a**) Parameters of the membrane shape: $H$ is the height, $r_b$ is the base edge radius, $r_p$ is the pore radius, $\phi$ and $\phi_{max}$ are the tangent angle of the membrane profile and is its maximal value, $r_{plug}$ is the radius of a circular membrane area to which the pulling force, $f_{pull}$, is applied.

(**b**) The energy landscape representing the system elastic energy normalized by the bending rigidity, $\frac{F_{el}}{\kappa}$, as a function of the normalized geometrical parameters, $\frac{H}{r_i}$ and $\frac{r_b}{ri}$.

(**c**) Shape diagram representing in terms of the parameters $\frac{H}{r_i}$ and $\frac{r_b}{r_i}$, where $r_i = \sqrt{\frac{\kappa}{2\gamma_0}}$ is the internal length scale, $\kappa$ is the membrane bending modulus, $\gamma_0$ is the membrane tension. The diagram represents the maximal tangent angle, $\phi_{max}$, which is colored coded and spans the range from low angles for flat shapes, through values close to $\frac{\pi}{2}$ for tubular shapes, and up to angles higher than $\frac{\pi}{2}$ for $\Omega$ shapes.

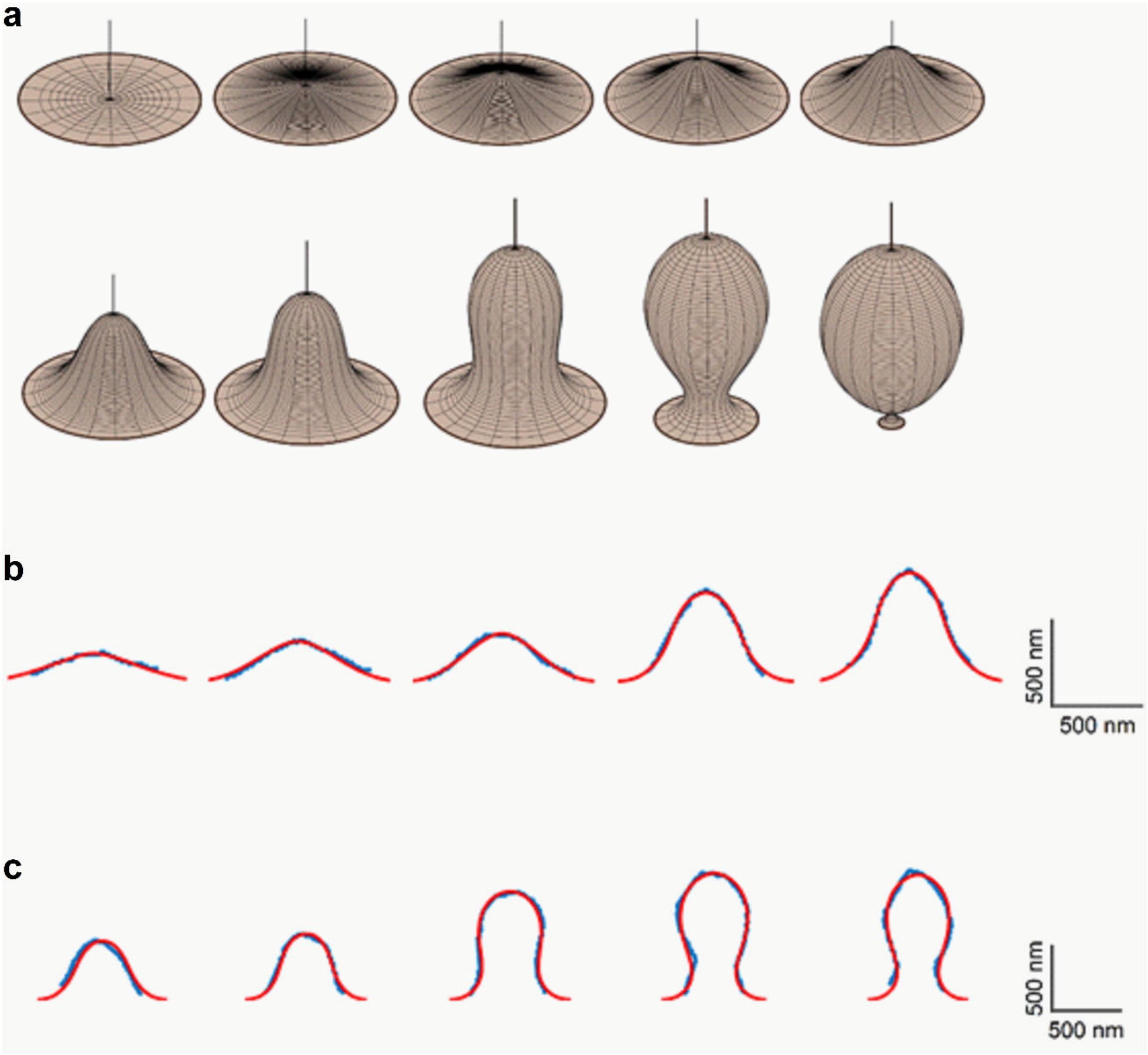


**Supplementary Figure 7. Evolution of bud shape.**

(**a**) Three-dimensional view of the computed membrane configurations representing the typical sequence of events: upper panel, Λ shapes with increasing height; lower panel, transformation of Λ-to Ω- shapes.

(**b-c**) Fitting of the computed shape profiles to the time sequence of shapes observed experimentally. The experimentally observed shape profiles were processed into pixel series shown by blue squares. The fitted model profiles are presented by red curves. The overlap between the fitted profiles and the original experimental images are presented in the main text (Figs. 7e and 7f, respectively).

**b**, Flat→Λ transition, see also Movie S7 for the computed versus the experimentally observed shape changes.

**c**, Λ→Ω transition, see also Movie S8 for the computed versus the experimentally observed shape changes.

#### Movie captions

**Movie S1. Actin filament in the act of pulling membrane inward to form Λ.**

Horizontal length of the image frame is 1.7 μm. The event is the same as the one shown in Fig. 1f. Green: $PH_G$-labelled plasma membrane; red: Lifeact-labelled F-actin. The video is played at real-time.

**Movie S2. Dynamin in the act of constricting Λ's base and Ω's pore**

Horizontal length of the image frame is 1.7 μm. The event is the same as the one shown in Fig. 4c. Green: $PH_G$-labelled plasma membrane; red: dynamin 1-mTFP1 puncta. The video is played 2 times as fast as the real-time.

**Movie S3. Cryo-electron tomogram of dynamin 1 with DOPS vesicles**

3-dimensional views of dynamin 1 assembled around the DOPS vesicle. Data are the same as those shown in Fig. 5d.

**Movie S4. CryoET segmentation of dynamin 1 around DOPS vesicle**

Dynamin 1 (yellow) forms helices around the DOPS vesicle (grey) with a diameter of ~166 nm. Data are the same as those shown for Fig. 5e, upper row.

**Movie S5. Cryo-ET segmentation of dynamin 1 around DOPS vesicle**

Dynamin 1 (yellow) forms helices around the DOPS vesicles (grey) with a diameter of ~85 nm. Data are the same as those shown for Fig. 5e, lower row.

**Movie S6. Simulated shape evolution for the Flat→Λ→Ω transition**

Evolution of simulated 3-D shapes for Flat→Λ (Fig. 7c) and Λ→Ω (Fig. 7d) transitions combined together, viewed from a different angle. The values of the pulling force, $f_{pull}$, and the base radius, $r_b$, are shown for each frame. All lengths are scaled by the intrinsic length, $r_i$.

Increasing the pulling force, $f_{pull}$, while constraining the base to a constant value, $r_b = 2.5 \cdot r_i$, results in Flat to Λ transition, which corresponds to the curve shown in (Fig. 7c). The shape transformation continues smoothly from Λ to Ω–shape by the base constriction upon conservation of the pulling force ($f_{pull} \cong 7.2 - 7.3\ f_i$) and membrane area, which corresponds to the curve shown in (Fig. 7d).

**Movie S7. Flat→Λ transition: the observed versus fitted simulation profiles**

Left: real-time video of the experimentally observed membrane profile evolution (scale bar: 500 nm).

Right: corresponding sequence of model-derived profiles obtained by fitting of the model parameters (see theoretical mathematical modeling). The fitted pulling force, $f_{pull}$, is labelled in every video frame.

**Movie S8. Λ→Ω transition: the observed versus fitted simulation profiles**

Left: real-time video of the experimentally observed membrane profile evolution (scale bar: 500 nm).

Right: corresponding sequence of model-derived profiles obtained by fitting of the model parameters (see theoretical mathematical modeling). The fitted base diameter, $r_b$, is labelled in every video frame.
